# Accumulation of Minor Alleles of Common SNPs in Schizophrenia

**DOI:** 10.1101/106815

**Authors:** Pei He, Xiaoyun Lei, Dejian Yuan, Zuobin Zhu, Shi Huang

## Abstract

Schizophrenia is a common neuropsychiatric disorder with a lifetime risk of 1%. A number of large scale genome wide association studies have identified numerous individual risk single nucleotide polymorphisms (SNPs) whose precise roles in schizophrenia remain unknown. Accumulation of many of these risk alleles has been found to be a more important risk factor. Consistently, recent studies showed a role for enrichment of minor alleles (MAs) in complex diseases. Here we studied the role of MAs in general in schizophrenia using public datasets. Relative to matched controls, schizophrenia cases showed higher minor allele content (MAC), especially for the sporadic cases. By linkage analysis, we identified 82 419 SNPs that could be used to predict 2.2% schizophrenia cases with 100% certainty. Pathway enrichment analysis of these SNPs identified 17 pathways, 15 of which are known to be linked with Schizophrenia with the remaining 2 associated with other mental disorders. These results suggest a role for a collective effect of MAs in schizophrenia and provide a method to genetically screen for schizophrenia.

Abbreviations

MAs: minor alleles
MAC: minor allele content
MAF: minor allele frequency
AUC: under the curve
TPR: True positive rate

## 1 Introduction

Schizophrenia is one of the most frequent neuropsychiatric disorders with a lifetime risk of 1% in the general population (McGrath et al., 2008; McGrath, 2007). This disease is often chronic and places a great burden on family and society. It is characterized by the occurrence of delusions, hallucinations, disorganized speech and behavior, impaired cognition, and mood symptoms (van Os and Kapur, 2009). Data from twin, family, and adoption studies showed strong evidence that schizophrenia is predominantly a genetic disorder with high heritability (Sullivan et al., 2003).

The precise mode of Schizophrenia inheritance is unclear and risk prediction using known genetic components is presently unrealistic. Based on investigating familial syndromes with schizophrenia-like phenotypes, two rare variants have been identified as associated with schizophrenia: the 22q11 deletion (Ivanov et al., 2003; Karayiorgou et al., 1995; Sporn A Fau - Addington et al., 2004) and a 1:11translocation (Blackwood et al., 2001; Hodgkinson et al., 2004). With the advent of copy number variants (CNVs) microarray technology, an increasing number of large rare deletions have been detected in schizophrenia patients (Levinson et al., 2011; Moreno-De-Luca et al., 2010; Walsh et al., 2008). However, the effect size associated with common CNVs is smaller than initially estimated (Wray and Visscher, 2010). In addition, many candidate genes for schizophrenia have been found by Genome-wide association studies (GWAS) (O'Donovan et al., 2008; Schizophrenia Psychiatric Genome-Wide Association Study, 2011). However, these SNPs are at frequencies of 20–80% in the general population and only account for a minimal increase in risk (Mulle, 2012; Tiwari et al., 2010). It is likely that schizophrenia may be related to accumulation of many risk alleles at thousands of loci (International Schizophrenia et al., 2009).

An allele can belong to either the major or the minor allele according to its frequency in the population and the minor allele (MA) has frequency (MAF) < 0.5. Most known risk alleles are MAs (Park et al., 2011). Our previous studies have shown that the collective effects of MAs may play a role in numerous traits and diseases (Yuan et al., 2014; Zhu et al., 2015a; Zhu et al., 2015b). Specifically, enrichment of genome wide common SNPs or MAs is associated with Parkinson's disease (Zhu et al., 2015b) and lower reproductive fitness in *C.elegans* and yeasts (Zhu et al., 2015a). We here studied the role of genome wide MAs as a collective whole in schizophrenia using previously published GWAS datasets.

## 2 Materials and Methods

### 2.1 Subjects

Two GWAS datasets of Cases and controls (phs000021.v3.p2, phs000167.v1.p1 (International Schizophrenia et al., 2009; O'Donovan et al., 2008; Stefansson et al., 2009; Suarez et al., 2006) were downloaded from database of Genotypes and Phenotypes (dbGaP). All subjects we selected for analysis are European-American ancestry population. The SNPs of all subjects were genotyped using AFFY_6.0 in genome-wide. Principal component analysis (PCA) using the GCTA tool was performed to analyze the genetic homogeneity of the subjects (Yang et al., 2011). Outliers were excluded through selection of the principal component values (Supplementary Table S1). Duplicated subjects were excluded, and the parents of cases were also excluded.

### 2.2 SNPs selection

All SNPs for analysis in this study are autosomal SNPs. In addition, we excluded SNPs showing departure from the Hardy-Weinberg equilibrium (P < 0.01), with missing data < 5%, and with MAF< 10^-4^. After these filters, there were 512 673 SNPs remaining (Table 1).

**Table 1.**
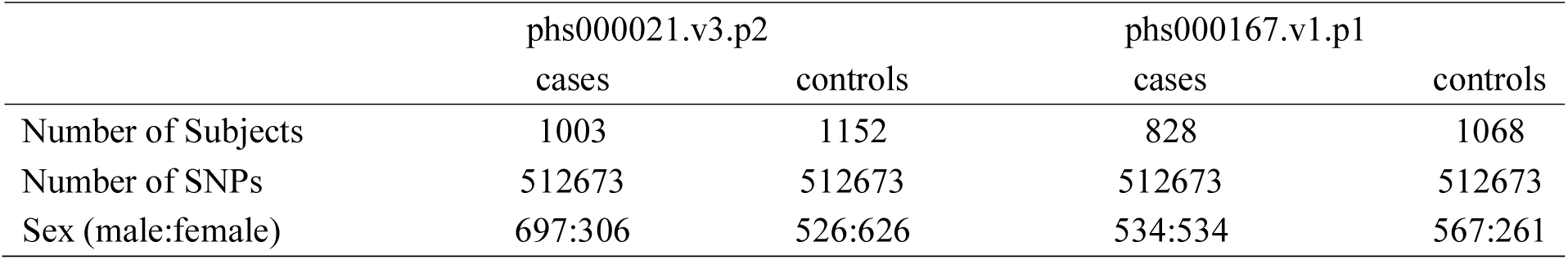
Description of datasets used in this study.

### 2.3 Statistical analysis

The Hardy-Weinberg equilibrium, missing data, MAF and logistic regression analysis were performed using PLINK Tools (Purcell et al., 2007). MAC per subject means the ratio of the total number of MAs divided by the total number of SNPs scanned (non-informative NN SNPs were excluded). The script for MAC calculation was previously described (Zhu et al., 2015b). Risk coefficient of each SNP was calculated with logistic regression test (equal to coefficient logistic regression test). The weighted risk score of a MA was calculated as follows: for homozygous MA, the risk coefficient was 1 x the coefficient, for heterozygous MA, it was 0.5 x the coefficient, for homozygous major allele, the coefficient was 0. The total weighted risk score from all MAs in a subject was obtained by summing up the weighted risk coefficient of all MAs by the script as described previously (Zhu et al., 2015b).

### 2.4 Genetic risk models construction and evaluation

125 prediction models were obtained from different combinations of MAF and p-value (Zhu et al., 2015b). For external cross-validation, the phs000021.v3.p2 study was used as training dataset, and the phs000167.v1.p1 study as validation dataset. Receiver operating characteristic (ROC) curves were used to describe the ability to differentiate cases and controls. True positive rate (TPR) is the proportion of cases with weighted risk scores higher than all of the controls. Area under the curve (AUC) and the TPR were calculated for each model by prism5.

For the internal cross-validation, a 10 fold cross-validation was used to test the models with good performance in external cross-validation. The models with TPR > 2% and AUC > 0.58 were chosen for internal cross-validation. Subjects in phs000021.v3.p2 were divided into 10 sub-sets randomly. When a sub-set was used as the validation data, the other 9 sub-sets were used as the training data. The cross-validation process was repeated 10 times, and the mean AUC and TPR values were calculated from these 10 results.

### 2.5 SNPs annotation and functional enrichment analysis

ANNOVAR (http://annovar.openbioinformatics.org/) was used to annotate SNPs (Wang et al., 2010). For functional enrichment analysis, WebGestalt (http://bioinfo.vanderbilt.edu/webgestalt/) tools were used for gene ontology annotation and pathway analysis according to Kyoto Encyclopedia of Genes and Genes (KEGG) (http://www.genome.jp/kegg/) (Wang et al., 2013; Zhang et al., 2005).

## 3 Results

### 3.1 Collective effects of minor alleles in Schizophrenia

We made use of the published GWAS datasets (phs000021.v3.p2 and phs000167.v1.p1). We first cleaned these datasets by removing outliers in principle component analysis (PCA) plots (see Figure in Supplementary Table S1). The cleaned datasets contains 1 003 cases and 1 152 controls in phs000021.v3.p2 dataset, and 828 cases and 1 068 controls in phs000167.v1.p1 dataset (Table 1). MA status of each SNP was then obtained by using the control cohort using MAF < 0.5 as cutoff. MAC of each subject was calculated, and the mean MAC of cases and controls was compared. The results showed that the mean MAC of schizophrenia cases is significantly higher than that of controls in both the phs000021.v3.p2 data (mean MAC [mean ± stdev], cases vs controls is 0.2235 ± 0.0010 vs 0.2233 ± 0.0011, *P* = 2.71E-06, t test) and the phs000167.v1.p1 data (cases 0.2251± 0.0011 vs controls 0.2249 ± 0.0011, *P* = 9.21E-04, t test, Supplementary Table S2). MAC values of both cases and controls showed normal distribution but cases were shifted slightly to the right or higher MAC values (Figure 1A and B).

**Figure 1.**
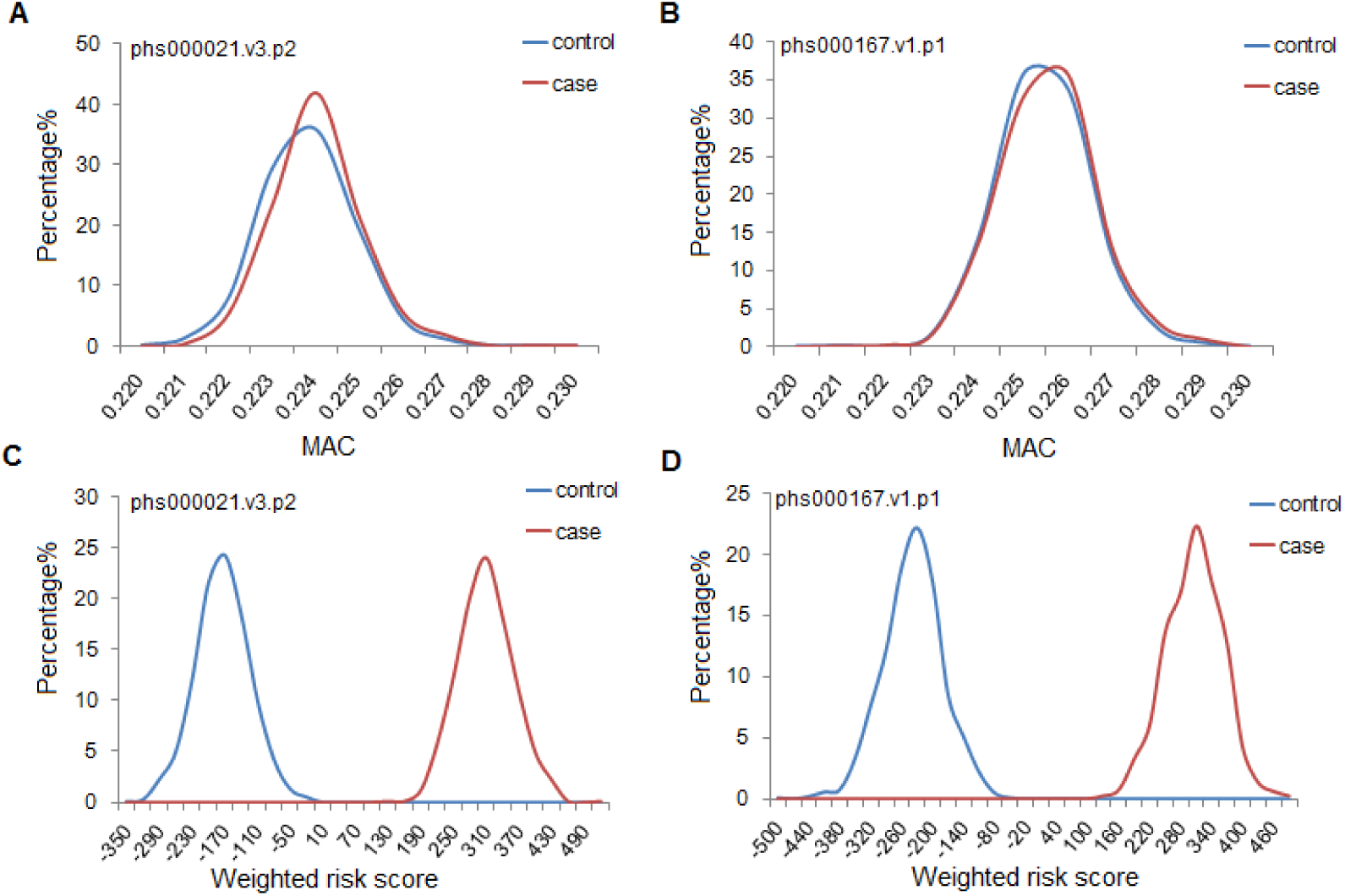
Minor allele distribution in cases and controls. Distribution of MAC (A, B) and weighted risk score (C, D) of case and control subjects in two different datasets. MAC: Minor allele content of SNPs with MAF < 0.5.

To study of the role of MAC in sporadic versus familial cases of schizophrenia, we further combined the subjects in the two datasets and calculated the MAC of each subject again. Then we compared the mean MAC of sporadic schizophrenia cases (n = 1217) to that of cases with family history of a psychotic illness (n = 493). We also compared these two groups of cases with the controls (n = 2220). The results showed that the mean MAC of sporadic schizophrenia cases (cases1) was slightly higher than cases with family history (cases2) (cases1 0.221505 ± 0.001039 vs cases2 0.221503 ± 0.001044，*P* = 0.49, one-way ANOVA). The MAC difference between sporadic cases and controls was significant (cases1 0.221505 ± 0.001039 vs controls 0.221412 ± 0.001054, *P* = 6.4E-03, one-way ANOVA), and more so than that between cases with family history and controls (cases2 0.221503 ± 0.001044 vs controls 0.221412 ± 0.001054, *P* = 0.04, one-way ANOVA, Supplementary Table S2). The results confirmed the expectation that collective effects of minor alleles should play more important role in sporadic Schizophrenia cases because familial cases may involve major effect mutations in a small number of genes.

We also calculated a risk coefficient score for each SNP by logistic regression analysis and obtained a weighted risk score based on the MA status and the risk coefficient score as previously described (Zhu et al., 2015b). The MAC of each individual was then converted into a weighted risk score by summing up the weighted risk scores of each SNP. The mean weighted risk score of cases was found to be far greater than that of controls in both datasets (cases 290.43 ± 55.86 vs controls -253.59 ± 57.90 in phs000021.v3.p2, *P* = ~0; cases 294.37 ± 50.77 vs controls -188.19 ± 50.50 in phs000167.v1.p1, *P* = ~0, t-test, Supplementary Table S2). This was apparent on a distribution plot of the weighted risk score with clearly separated cases and controls (Figure 1C and D).

### 3.2 Evaluation of risk prediction models

In order to get an optimal MAs model or a subset of MAs for risk prediction, we divided the MAs into 5 groups according to MAF (<0.5, <0.4, <0.3, <0.2, and <0.1, Fig 2). We performed logistic regression analysis and obtained the p-values for each SNP. Based on these p-values, we divided each group into 25 subgroups and obtained a of 125 prediction models (Fig 2, Supplementary Table 3). We then performed external cross-validation and internal cross-validation analyses to test these models. In external cross-validation, we used phs000021.v3.p2 as the training dataset and phs000167.v1.p1 as the validation dataset. We then used the receiver operator characteristic (ROC) curve to examine the discriminatory capability or area under the curve (AUC) of each model in the testing dataset. We found 17 models with AUC > 0.58 and true positive rate (TPR) > 2%. Among these models, the best TPR is 2.78%, and the best AUC is 0.6 (Fig 2 and Supplementary Table S3).

**Figure 2.**
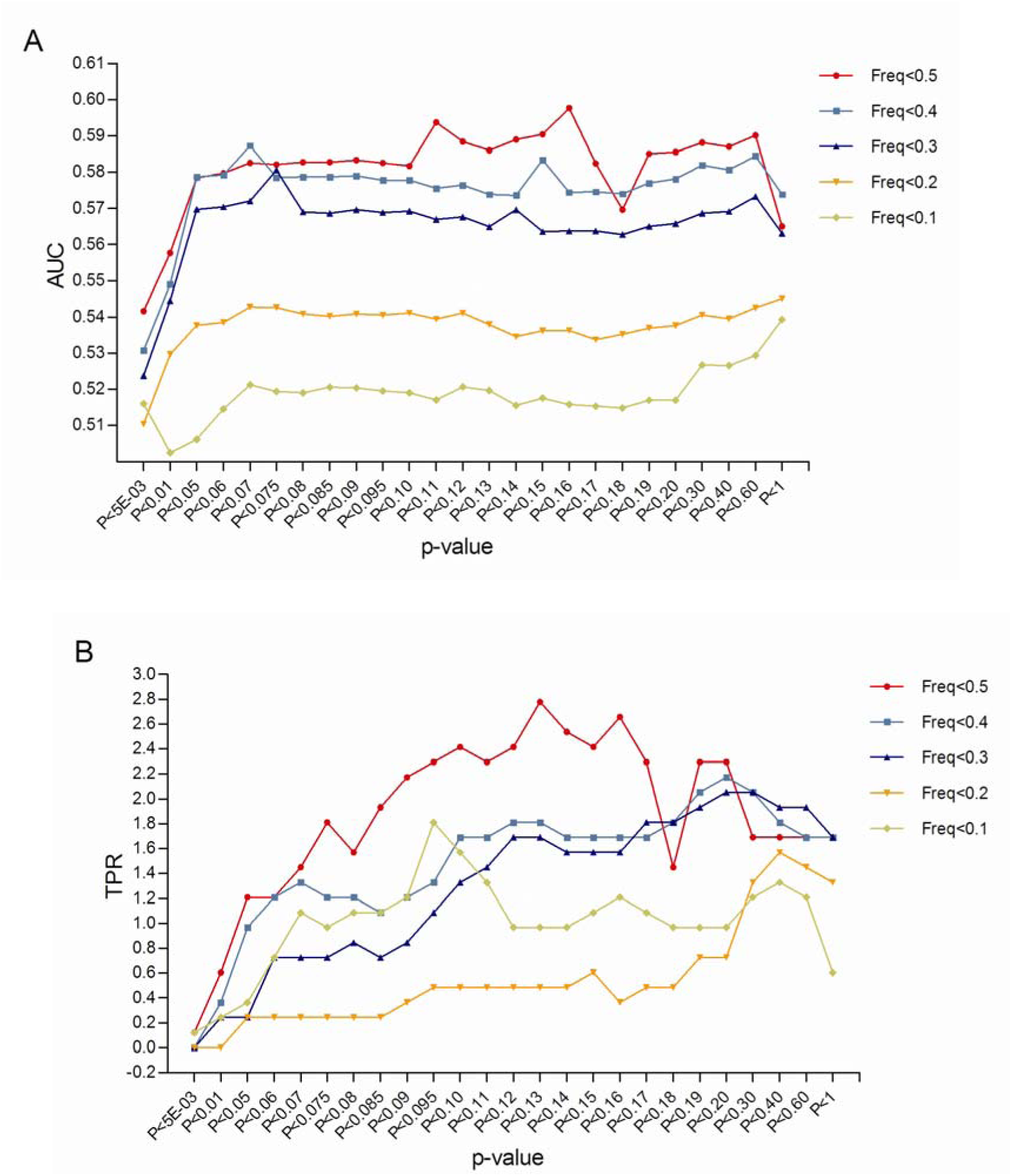
Discriminatory ability of different prediction models. SNPs were divided into 5 groups based on MAF, each group was further divided into 25 subgroups based on p-values from the logistic regression test and 125 prediction models were obtained. AUC (A) and TPR (B) were calculated using a training dataset and a validation dataset to evaluate the discriminatory ability.

A 10 fold internal cross-validation analysis with these 17 models was further performed using phs000021.v3.p2 dataset. Each model was analyzed 10 times, and the mean AUC and TPR were calculated. The best model had AUC 0.62 (95%CI, 0.5919-0.6367) and TPR 2.2% (95%CI, 0.8786%-3.4957%) in internal cross-validation analysis, and AUC 0.6 (95%CI, 0.5678-0.6374) and TPR 2.67% (95%CI, 1.672%-3.995%) in external cross-validation analysis. There were 82 419 SNPs in this model with MAF < 0.5, and each MA had a p-value < 0.16 (Figure 2 and Supplementary Table S3).

We next tested whether the set of 82 419 SNPs is relatively specific to schizophrenia. We compared these SNPs with the previously identified 37 564 SNPs specific for Parkinson’s disease (Zhu et al., 2015b). Only 1 239 SNPs were found shared between these two sets, indicating that different diseases may be linked with different sets of SNPs.

### 3.3 SNPs annotation

We next examined the potential functions of the 82 419 SNPs by annotating them with the ANNOVAR software (Wang et al., 2010). There were 82 834 SNPs annotation results in total due to the fact that some SNPs may lie in between two genes and could hence generate two annotation results. We found 0.956% of SNPs in exonic regions (Table 2, Supplementary Table S4).

**Table 2.** Statistics of SNPs distribution.

| **Locus** | **Number of SNPs** | **Percentage%** |
| --- | --- | --- |
| Exonic | 792 | 0.956 |
| Intergenic | 44440 | 53.649 |
| Intronic | 30742 | 37.113 |
| UTR3 | 821 | 0.991 |
| UTR5 | 80 | 0.097 |
| Upstream | 357 | 0.431 |
| Downstream | 536 | 0.647 |
| upstream,downstream | 8 | 0.010 |
| Splicing | 4 | 0.005 |
| ncRNA-intronic | 4735 | 5.716 |
| ncRNA-exonic | 319 | 0.385 |

We mapped the 82 419 SNPs to gene loci using WebGestalt tools, and found 6 588 genes (Supplementary Table S4). These genes were characterized using Gene Ontology in WebGestalt according to biological process, molecular function, and cellular component. As shown in Table 3, most of these genes were related to cytoskeletal proteins, phospholipid, anion and actin biding, GTPase regulators, transmembrane receptor protein tyrosine kinases, transmembrane receptor protein kinases, small GTPase regulators, and mental ion transmembrane transporter activities, and nucleoside-triphosphatase regulators.

**Table 3.**
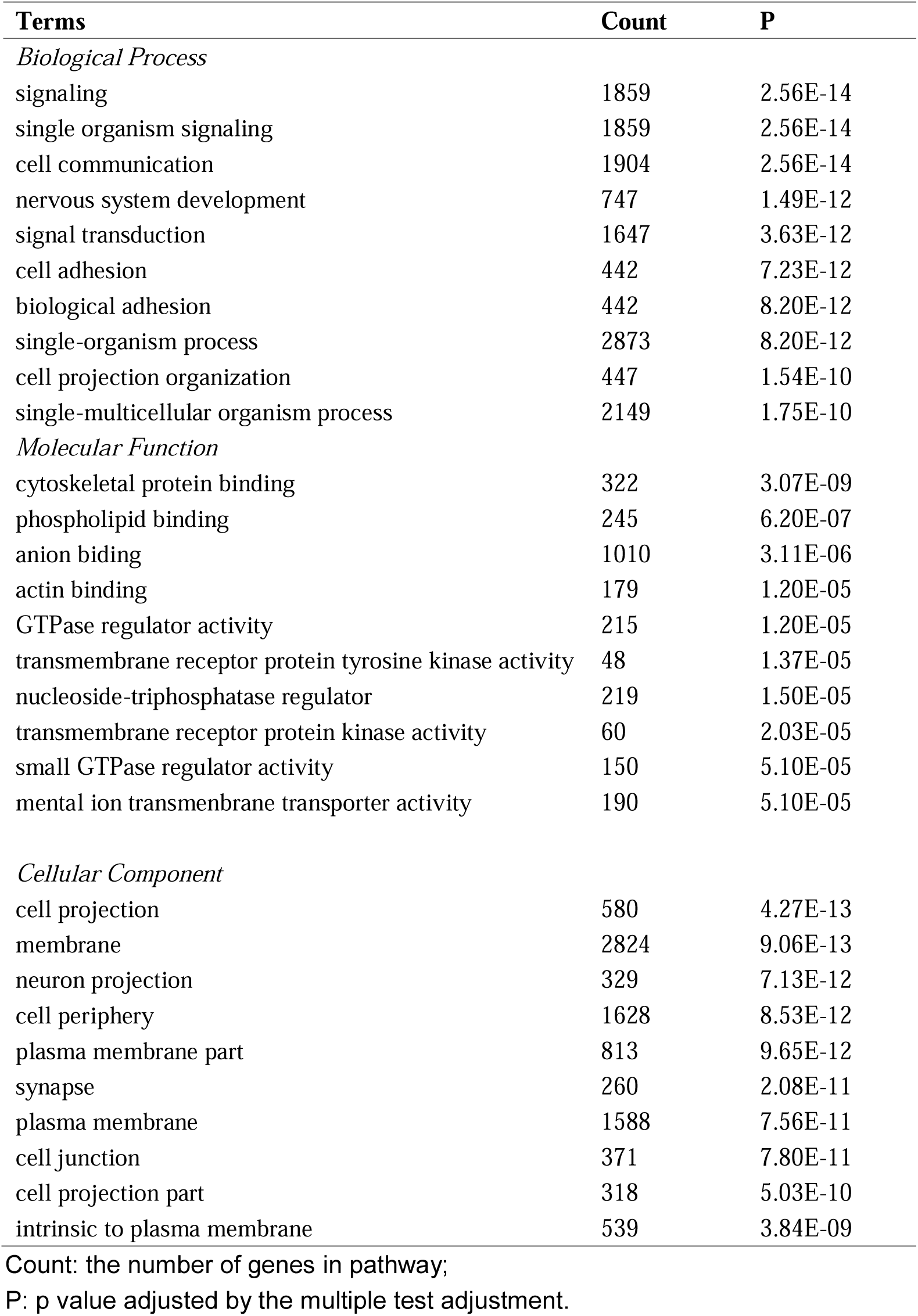
Categories of top-10 significantly enriched genes.

Pathway analysis was carried out on these 6 588 genes according to the Kyoto Encyclopedia of Genes and Genomes (KEGG) using WebGestalt tools. A total of 17 pathways were identified with *P* < 0.05 (after multiple test adjustment) (Table 4, and Supplementary Table S5). All of these signaling pathways have been shown to have a role in schizophrenia except the gastric acid secretion and bile secretion pathway that may play a role in autism, depression (Horvath et al., 1999; Padol et al., 2012), and Alzheimer's disease (Simpson et al., 1994; Winkler et al., 2015). The pathways linked with schizophrenia include focal adhesion(Fan et al., 2013), axon guidance (Chen et al., 2011), calcium signaling pathway (Berridge, 2013; Hertzberg et al., 2015; Lidow, 2003), ECM-receptor interaction (Lubbers et al., 2014), vascular smooth muscle contraction (Sakakibara et al., 2012), arrhythmogenic right ventricular cardiomyopathy (Kawasaki et al., 2015), regulation of actin cytoskeleton (Criscuolo and Balledux, 1996; Zhao et al., 2015), long-term potentiation (Frantseva et al., 2008; Hasan et al., 2011; Salavati et al., 2015), MAPK signaling pathway (Funk et al., 2012), ABC transporters (Akamine et al., 2016), neuroactive ligand-receptor interaction (Adkins et al., 2012), GnRH signaling pathway (Brambilla F Fau - Rovere et al., 1976), salivary secretion (Toone Bk Fau - Lader and Lader, 1979), cell adhesion molecules (CAMs) (Webster et al., 1999; Zhang et al., 2015) and dilated cardiomyopathy (Finsterer and Stollberger, 2016; Volkov Vs Fau - Volkov and Volkov, 2013).

**Table 4.** Significantly enriched KEGG pathways from WebGestalt Toolkit.

| **KEGG pathway** | **O** | **E** | **R** | **P** |
| --- | --- | --- | --- | --- |
| Focal adhesion | 105 | 76.3 | 1.38 | 0.0027 |
| Axon guidance | 70 | 47.84 | 1.46 | 0.0027 |
| Calcium signaling pathway | 86 | 62.27 | 1.38 | 0.0049 |
| ECM-receptor interaction | 47 | 31.76 | 1.48 | 0.0112 |
| Vascular smooth muscle contraction | 58 | 40.42 | 1.44 | 0.0112 |
| Arrhythmogenic right ventricular cardiomyopathy (ARVC) | 43 | 28.46 | 1.51 | 0.0112 |
| Gastric acid secretion | 39 | 25.98 | 1.5 | 0.0173 |
| Regulation of actin cytoskeleton | 99 | 76.71 | 1.29 | 0.0173 |
| Long-term potentiation | 38 | 25.16 | 1.51 | 0.0173 |
| MAPK signaling pathway | 118 | 94.03 | 1.25 | 0.0178 |
| ABC transporters | 27 | 16.91 | 1.60 | 0.0264 |
| Neuroactive ligand-receptor interaction | 114 | 91.97 | 1.24 | 0.0316 |
| GnRH signaling pathway | 48 | 34.64 | 1.39 | 0.0395 |
| Salivary secretion | 39 | 27.22 | 1.43 | 0.0398 |
| Cell adhesion molecules (CAMs) | 61 | 46.19 | 1.32 | 0.0461 |
| Dilated cardiomyopathy | 45 | 32.58 | 1.38 | 0.0474 |
| Bile secretion | 38 | 26.81 | 1.42 | 0.0485 |

O: the number of genes in our gene set and also in the category; E: the expected gene number in the category; R: ratio of enrichment; P: p value adjusted by the multiple test adjustment.

## 4 Discussion

In this study, we showed enrichment of MAs in schizophrenia cases relative to matched controls. We also identified a set of 82 419 SNPs that can predict with certainty a fraction of schizophrenia cases. These results are consistent with previous work on other complex diseases and traits (Yuan et al., 2014; Zhu et al., 2015a; Zhu et al., 2015b).

There were reports of male bias in schizophrenia (Aleman et al., 2003). The ratio of males to females in cases of phs000021.v3.p2 data was 2.28 but was close to 1 in the other dataset. We however did not observe significant differences in MAC values between male and female cases in both datasets. Thus, MAC may play a similar role in both sexes.

Recent studies have shown that a much larger than expected portion of the human genome may be functional (Fung et al., 2014; Hu et al., 2013; Sauna and Kimchi-Sarfaty, 2011). Genetic diversities are at saturation levels as indicated by the observation that higher fractions of fast evolving SNPs are shared between different human groups (Yuan et al., 2017). This raises the question of what selection forces are keeping genetic diversity levels from increasing with time. By linking the total amount of SNPs or MAs in an individual to complex diseases and traits, it is clear that complex diseases could serve as a negative selection mechanism to prevent abnormal increase in SNP numbers in an individual. It is intuitively obvious that the overall property of the genome as a whole should be linked with the wellbeing of an organism. Our results here on schizophrenia further confirmed the hypothesis we put forward before that a highly complex and ordered system such as the human brain must have an optimum limit on the level of randomness or entropy in its building parts or DNAs (Zhu et al., 2015b).

It has been difficult to use any genetic markers or combinations of them to predict risk of schizophrenia. We here identified a set of 82 419 SNPs that could predict 2.2% cases with 100% specificity. Although this is still a low percentage, it may still prove valuable for prenatal diagnosis of schizophrenia. The set of 82 419 SNPs specific for schizophrenia was highly linked with pathways known to be involved in the disease, thereby validating our method of looking for disease specific set of SNPs. This set is much larger than any known from previous studies (International Schizophrenia et al., 2009). This large collection of risk alleles is not unexpected if most genome sequences are functional as explained by the maximum genetic diversity (MGD) theory (Huang, 2008; Huang, 2009; Huang, 2016), which inspired this work in the first place. Future studies using larger sample sizes may help identify a more specific set of risk SNPs that could predict higher fraction of cases.

## Contributors

Shi Huang and Zuobin Zhu designed the study and wrote the protocol. Pei He, Xiaoyun Lei and Dejian Yuan managed the literature searches and analyses. Pei He undertook the statistical analysis and wrote the first draft of the manuscript. All authors contributed to and have approved the final manuscript.

## Conflict of interest

The authors declare that they have no conflict of interest.

## Role of the funding source

This work was supported by the National Natural Science Foundation of China grant 81171880 and the National Basic Research Program of China grant 2011CB51001 (S.H.).

## Acknowledgments

We thank NINDS dbGaP GWAS Data Repository, the datasets (phs000021.v3.p2, phs000167.v1.p1) necessary for our analysis were obtained from it; we also thank all the Contributing Investigator of raw data.

